# Immediate early proteins of herpes simplex virus transiently repress viral transcription before subsequent activation

**DOI:** 10.1101/2022.06.13.495981

**Authors:** Laura E.M. Dunn, Claire H. Birkenheuer, Rachel Dufour, Joel D. Baines

## Abstract

Herpes simplex virus 1 (HSV-1) utilizes cellular RNA polymerase II (Pol) to transcribe its genes in one of two phases. In the latent phase, viral transcription is highly restricted but during the productive lytic phase, more than 80 genes are expressed in a temporally coordinated cascade. In this study, we used precision nuclear Run On followed by deep Sequencing (PRO-Seq) to characterize early viral transcriptional events using HSV-1 immediate early (IE) gene mutants, corresponding genetically repaired viruses, and wild type virus. Unexpectedly, in the absence of the IE genes ICP4, ICP22 or ICP0 at 1.5 hpi we observed high levels of aberrant transcriptional activity across the mutant viral genomes, but substantially less on either wild type or the congenic repaired virus genomes. This feature was particularly prominent in the absence of ICP4 expression. Cycloheximide treatment during infection with both the ICP4 and ICP22 mutants and their respective genetic repairs did not alter the relative distribution of Pol activity, but increased overall activity across both viral genomes, indicating that both virion components and at least some *de novo* protein synthesis were required for full repression. Overall, these data reveal that prior to their role in transcriptional activation, IE gene products and virion components first repress transcription and that the HSV-1 lytic transcriptional cascade is mediated through subsequent de-repression steps.

**Importance:** Herpes simplex virus 1 (HSV-1) transcription during productive replication is believed to comprise a series of activation steps leading to a specific sequence of gene expression. Here we show that virion components and immediate early (IE) gene products ICP0, ICP4 and ICP22 first repress viral gene transcription to varying degrees before subsequently activating specific gene subsets. It follows that the entire HSV transcriptional program involves a series of steps to sequentially reverse this repression. This previously uncharacterized repressive activity of IE genes very early in infection may represent an important checkpoint allowing HSV-1 to orchestrate either the robust lytic transcriptional cascade or the more restricted transcriptional program during latency.

## Introduction

Herpes simplex virus 1 (HSV-1) expresses its double-stranded DNA genome using cellular RNA polymerase II (Pol) in two phases. During the latent phase in which the genome is maintained as an episome in sensory neurons, viral transcription is mostly limited to the production of the latency transcript (LAT). In the productive phase which usually occurs in epithelial cells, transcription of its more than 80 genes occurs in a cascade involving sequential activation of different gene subsets: from pre-immediate early, followed sequentially by immediate early (IE, or α), early (E or β), leaky late (LL or γ_1_), and true late (L, γ_2_) genes (1,2). To initiate productive replication, HSV commandeers Pol for viral transcription by introducing VP16 from the virion into the newly infected cell. VP16 binds a specific motif in viral IE promoters and recruits cellular transcription factors, HCF and Oct-1, to these sites to drive Pol initiation (3,4). Similar to cellular genes, viral gene expression is regulated through promoter proximal pausing (PPP) of Pol, followed by release into the gene body (5).

The IE genes are expressed by 1 hour post-infection and include α0, α4 and α22/US1, encoding ICP0, ICP4 and ICP22, respectively (6). ICP0 and ICP4 are also virion components that are introduced into the cell upon the initiation of infection (7). ICP4 is an essential activator and repressor of both viral (8) (9) and host genes(10), while ICP0 is an E3 ubiquitin ligase that leads to degradation of a set of cellular proteins that would otherwise silence the viral genome early in infection (11). ICP22 negatively regulates Pol processivity and release to elongation through reduction of serine 2 phosphorylation on the Pol C-terminal domain (CTD) (12). This negative elongation activity of ICP22 is important for enhancing promoter proximal pausing (PPP) on viral IE genes, and controlling anti-sense transcription on the compact HSV-1 genome (13).

The current studies were undertaken to further clarify the roles of ICP4, ICP0, and ICP22 in early viral transcription. We used Precision nuclear Run On followed by deep Sequencing (PRO-Seq) to identify sites of Pol activity to base-pair resolution on the genomes of viruses bearing mutations in α0, α4 or α22, and corresponding viruses in which these loci were genetically restored.

## Results

### In the absence of ICP4, Pol activity is dysregulated throughout HSV-1 infection

We first used the ICP4 truncation mutant n12, derived from the KOS HSV-1 strain (1) to investigate the role of ICP4 during transcription. To allow comparison with virus that was otherwise genetically identical to the mutant, a repair virus was generated from n12 by recombination in infected cells with wild type α4 DNA. Expression of full length ICP4 was confirmed by immunoblotting (Fig. S1A). A single-step growth curve of the repair compared to n12, and WT HSV-1(KOS) showed restored replication kinetics and high final yields relative to n12, with only slightly reduced replication compared to WT (Fig. S1B, C). HEp-2 cells were then infected at a multiplicity of infection (MOI) of 5.0 with either n12 or repaired virus, nuclei harvested at 1.5, 3 or 6 hours post infection (hpi), and PRO-Seq performed.

HEp-2 cells harvested at 3 and 6 hpi were chosen for comparison to previous HSV-1 PRO-Seq studies (5,13). We included the 1.5 hpi time point to measure transcription prior to the onset of viral DNA replication (5).

Visualization of the data on a genome browser revealed substantial differences in the pattern of Pol activity on the n12 genome compared to the repair genome (Fig. 1A, first panel). Consistent with the strict temporal expression of herpesvirus genes, Pol activity on the repair virus genome was restricted to IE genes at 1.5 hpi. In contrast, strong Pol activity was distributed across the entire n12 genome. At 3 hpi, the distribution of Pol activity on the two viral genomes was more similar, but activity on the pre-IE gene L/ST, the IE genes UL54 (α27), α4, α0, US1 (α22), US12 (Fig. 1A, second panel) and many late genes remained higher in n12 than the repair. By 6 hpi, intense Pol activity of n12 was limited almost entirely to IE genes (and UL39), while activity on the repair virus genome was more evenly distributed among genes of all temporal classes (Fig. 1A, third panel).

**Figure 1:**
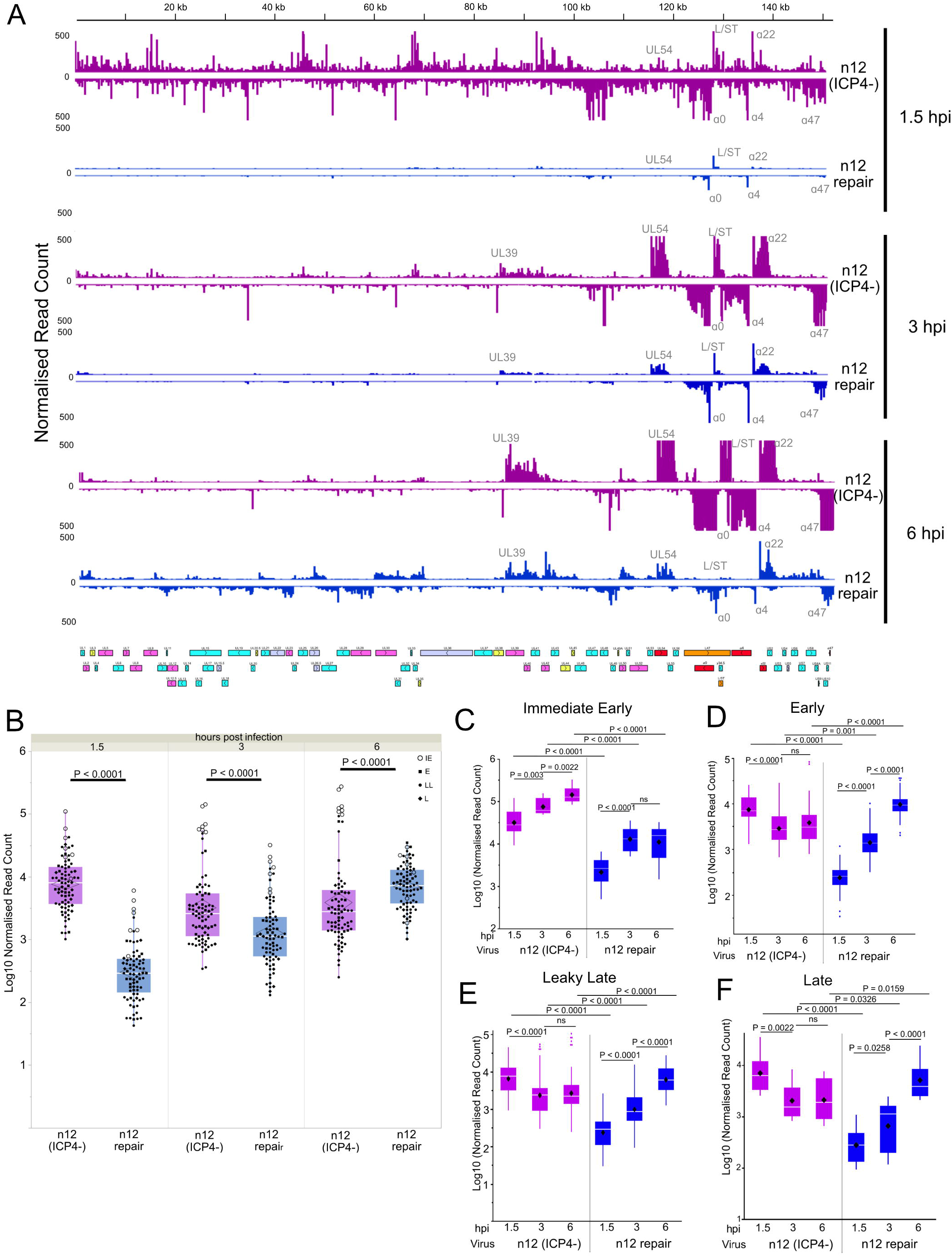
Dysregulated Pol activity throughout HSV-1 infection in the absence of ICP4. HEp-2 cells were infected with either the ICP4-mutant, n12, or its genetically restored repair and PRO-Seq performed at 1.5, 3 and 6 hpi. **(A)** Genome browser view of distribution of PRO-Seq reads (mean of 2 biological replicates, normalized to drosophila spike-in) across the HSV-1 genome. External repeat sequences were deleted for the sequencing alignment. IE gene peaks are noted. **(B)** Log10 normalized read counts of all HSV-1 genes. Mann-Whitney U test was used to estimate statistical significance. Log10 normalized read counts of **(C)** Immediate Early, **(D)** Early, **(E)** Leaky Late and **(F)** Late genes. Black diamonds indicate mean. Kruskal-Wallis test was performed to estimate statistical significance. ns = P > 0.05.

Quantification of sequencing reads aligning to individual HSV-1 genes confirmed a significant increase in Pol activity in n12 over the repair virus at 1.5 and 3 hpi (Fig. 1B). At 6 hpi the total viral reads remained higher in n12, but this was due solely to intense IE gene activity (Figs. 1C) that did not occur in repair virus. Over time, activity on later gene classes of n12 (Fig. 1D, E, F) decreased, while in repair infection Pol activity increased on all gene classes, except for IE genes between 3-6 hpi (Fig. 1C). Overall, these data indicate that ICP4 decreases Pol activity on all viral genes immediately after infection, and increases E, LL and L activity later in infection.

PRO-Seq profiles of Pol occupancy on most HSV-1 genes follow that of cellular genes, with peaks at the 5’ PPP site, and at the 3’ cleavage/Poly(A) site (5). The promoters of the IE genes α4 and US1 showed clear PPP at all time points on the repair genome (Fig. 2A). High read counts on n12 IE genes obscured patterns of Pol activity on the same scale so the browser was adjusted to a comparable read count depth between the two viruses. This revealed strong PPP on α4 and US1 throughout infection in n12. Other transcriptionally active regions in n12 lacked apparent PPP at 1.5 hpi including E genes UL29 and UL30 (Fig. 2B), the LL genes UL18, UL19 and UL20 and the L genes UL20.5 and UL22 (Fig. 2C). The presence of a promoter peak was also assessed by examining the relative read density over a gene and using bootstrap confidence of fit to indicate statistical differences. To remove any uncertainty of whether a read was from a promoter region or gene body, only genes with no overlapping transcripts and with a defined TSS were included in the analysis. We also separately analysed genes that were robustly transcribed in repair infection at each time point as it was assumed that these gene were transcribed in a temporally correct fashion. The set of appropriate positively regulated genes were determined by having a mean read per bp ≥ 1 standard deviation of mean and was termed “repair-active”. Genes outside of this set (i.e., that were transcribed robustly in n12 but not repair at a given time point) comprised a “repair-repressed” subset.

**Figure 2:**
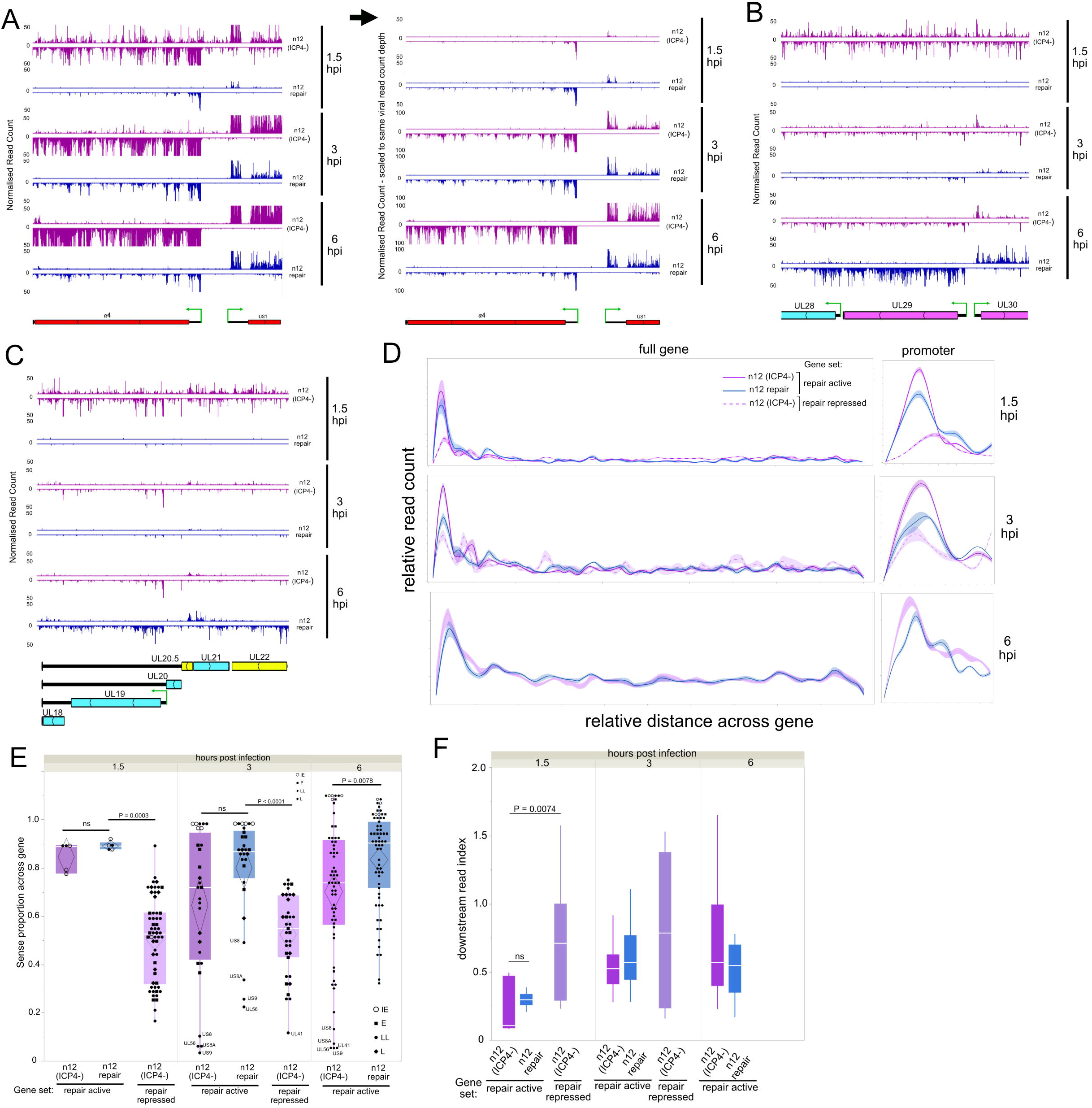
Aberrant transcription on the HSV-1 genome at 1.5 hpi in the absence of ICP4. HEp-2 cells were infected with either the ICP4-mutant, n12, or its genetically restored repair and PRO-Seq performed at 1.5, 3 and 6 hpi. High resolution genome browser view PRO-Seq tracks HSV-1 regions (mean of 2 biological replicates, normalized to drosophila spike-in): **(A)** α4 and US1, shown scaled to equal sequencing depth and equal viral read counts, **(B)** Ul28, UL29 and UL30, **(C)** UL18, UL19, UL20, UL20.5, UL21, UL22. IE genes: red, E genes: pink, LL genes: blue, L genes: yellow. **(D)** Spline interpolation analysis of the relative distribution of reads across repair-active genes and repair-repressed genes A closer view of the promoter region is also shown. The bootstrap confidence of fit is shown in the shaded area. Proportion of reads mapping to the sense strand on **(E)** all HSV-1 repair-active and repair-repressed genes. **(F)** Downstream-read index of repair-active and repair-repressed genes. Mann-Whitney U test was used to estimate statistical significance between viruses or gene sets at each time point. ns = P > 0.05. Repair-active: genes that are robustly transcribed in both repair and mutant. Repair-repressed: genes that are robustly transcribed only in mutant

Despite a large difference in total read counts on viral genes at 1.5 hpi, the distributions of reads across entire repair-active genes (i.e., relative read density) of n12 and repair were equivalent (Fig. 2D, top panel). Analysis of the promoter regions of active genes showed strong PPP on both viruses but with significantly higher PPP in n12 than in repair. A reduced but discernible promoter peak was visible on n12 repair-repressed genes. At 3 hpi a similar pattern was seen (Fig. 2D, middle panel). When all genes in repair were included in the repair-active subset due to robust transcription at 6 hpi, both viruses had a similar pattern of read density across individual genes. Pause indices confirmed this trend and revealed that the patterns of pausing changed similarly throughout infection (Fig. S2A-C). These data indicate that the ICP4-mediated repression of most genes at IE times is due to a reduction in total Pol occupancy across entire genes rather than an increase in Pol pausing.

PRO-Seq visualization indicated extensive Pol activity on the antisense strands of genes in n12 at 1.5 hpi. To examine the regulation of sense-to-antisense transcription, the proportion of sense reads was calculated on isolated genes that lacked genes on the opposite strand.

There was no significant difference in the levels of sense transcription between the two viruses on repair-active genes at 1.5 and 3 hpi, and both viruses had a high proportion of sense reads (Fig. 2E). However, the repair-repressed genes had a significant increase in antisense transcription at 1.5 and 3 hpi in n12. While n12 did have a significantly reduced global level of sense transcription at 6 hpi, this was largely due to high levels of antisense transcription on US8, US8A, US9, UL41 and UL56. Closer analysis of these regions revealed the presence of an upstream ICP4-independent promoter (US12, UL54, US1 and UL39) potentially driving read-through antisense Pol activity (Fig. S3A-D). This activity was also visible on the repair genome, indicating that it is a feature of HSV-1 transcription.

To determine whether detected intergenic Pol activity could be a result of read-through of a transcriptional termination signal (TTS) (AAUAA), a downstream-read index was calculated by dividing the reads per bp 150 bp downstream of a TTS, by the reads per bp in the upstream ORF. For reads to be accurately assigned, only singular unnested genes with a defined TTS more than 150bp away from a transcription start site (TSS) were included. At 1.5 hpi, there was no significant difference in the downstream index on repair-active genes between n12 and repair (Fig. 2F). However, there was a significantly higher proportion of downstream transcription on repair-repressed genes at 1.5 hpi. By 3 and 6 hpi, there were no significant differences in intergenic transcription on the gene sets between viruses, suggesting proper termination was restored at these time points even in the absence of ICP4.

Overall, the PRO-Seq data at 3 and 6 hpi was consistent with ICP4 mutant phenotypes, primarily consisting of a failure to repress IE genes and to promote transcription of later gene classes (14,15). However, the finding that ICP4 was required to repress Pol activity on virtually all viral genes at 1.5 hpi was unexpected. Pol activity on n12 IE genes was aberrantly high at 1.5 hpi, suggesting accelerated initiation. However, these genes displayed low levels of antisense and downstream transcription, and strong PPP peaks, suggesting pausing and termination approached normal levels. In contrast, Pol activity on genes of later kinetic classes was indiscriminate and included antisense and intergenic transcription.

Restoration of features consistent with proper termination and PPP returned in n12 by 3 and 6 hpi indicating that proteins other than ICP4 act to improve transcriptional control after 1.5 hpi. Similar results as above were obtained in PRO-Seq experiments performed at 1.5 hpi of primary human foreskin fibroblasts (HFF) showing that ICP4-mediated repression of viral transcription occurs in multiple cell types (Fig. S5A-F).

### Virion-associated and *de novo*-expressed ICP4 are required for full transcriptional repression at 1.5 hpi

We next asked whether the role of ICP4 to repress Pol early in infection required *de novo* protein synthesis or could be mediated by ICP4 entering the cell as a virion component. It should be noted that n12 virions do incorporate some ICP4 from the E5 cell line used to support n12 growth (16). HEp-2 cells were infected in the presence or absence of the protein synthesis inhibitor cycloheximide (CHX), followed by PRO-Seq at 1.5 hpi. Overall, CHX treatment led to a significant increase in reads aligning to viral genes on both the repair and n12 genome (Fig. 3A), indicating that *de novo* protein synthesis was required for full repression. However, Pol activity on the genes which reached the transcriptionally robust activity threshold (read per bp ≥ 1 standard deviation of mean) were unaffected by CHX treatment in both virus infections. Moreover, the proportion of sense transcription on the repair-active genes in repair was unchanged by CHX treatment and was not reduced to the lower proportion seen across n12 (Fig. 3B).

**Figure 3:**
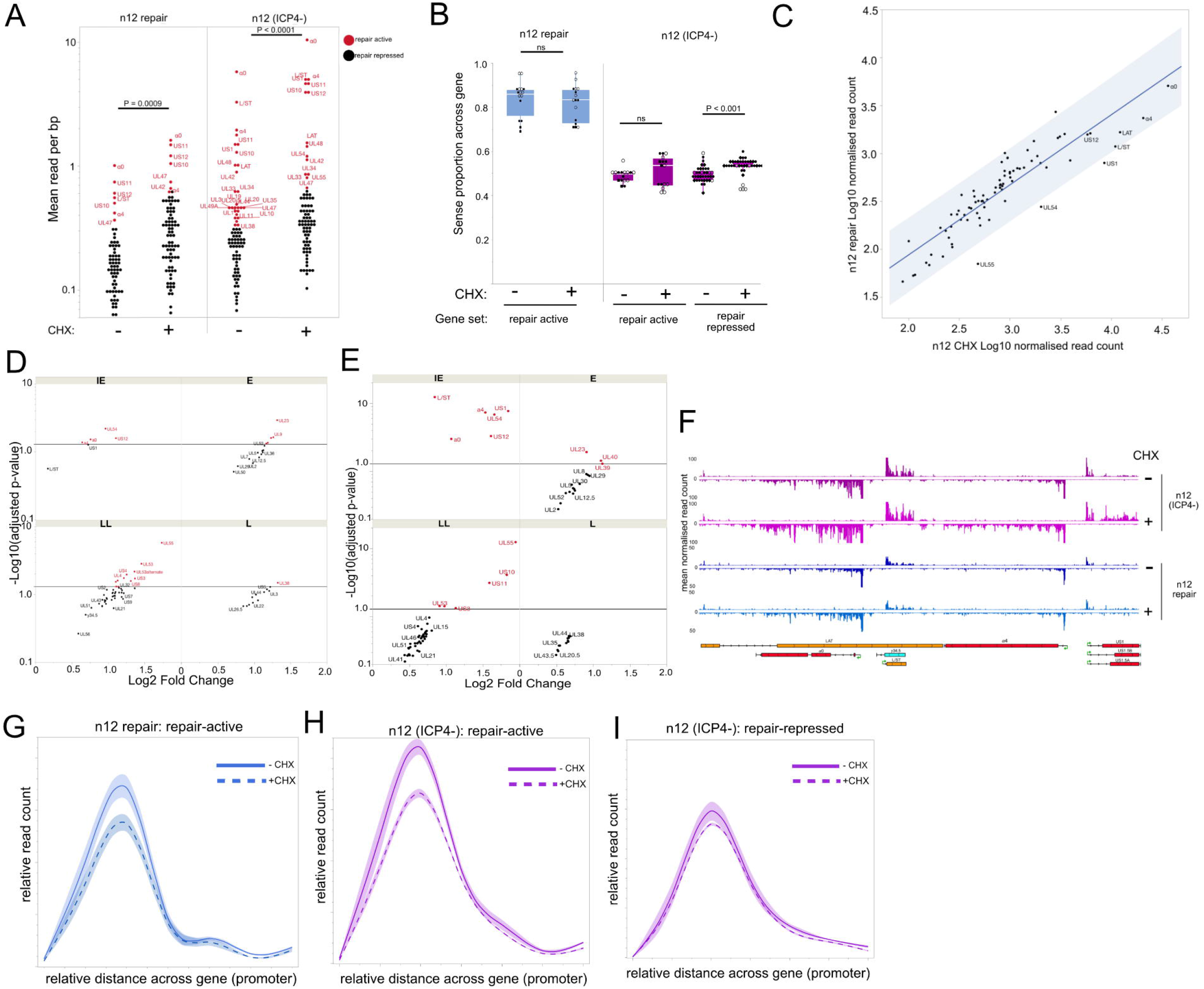
Both virion-associated and *de novo*-synthesized ICP4 are required for early transcriptional repression on the HSV-1 genome at 1.5 hpi. HEp-2 cells were infected with either the ICP4-mutant, n12, or its genetically restored repair in the presence or absence of the protein synthesis inhibitor cycloheximide (CHX) and PRO-Seq performed at 1.5 hpi. n=2 for each treatment, normalized to drosophila spike-in. **(A)** Mean read per bp across HSV-1 genes, red genes indicate those that were defined as being robustly transcribed. **(B)** Proportion of reads mapping to the sense strand on repair-active and repair-repressed genes. **(C)** Scatterplot of HSV-1 gene reads after CHX treatment in repair and n12 infection. Shaded area indicates confidence prediction (mean ±standard error). DeSeq2 Log2 fold change comparison of HSV-1 gene reads, separated by gene class of **(D)** repair CHX treated relative to repair untreated, **(E)** n12 CHX treated relative to n12 untreated. IE: immediate early, E: early, LL: leaky late, L: late. Genes with a fold change adjusted p-value ≤ 0.05 are shown in red. **(F)** PRO-Seq tracks of the HSV-1 α0-US1 IE gene region (mean of 2 biological replicates, normalized to drosophila spike-in). Spline interpolation analysis of the relative distribution of reads across promoter regions in n12 repair repair-active genes **(G)**, n12 repair-active genes **(H)** and n12 repair repressed **(I)**. The bootstrap confidence of fit is shown in the shaded area. Repair-active: genes that are robustly transcribed in both repair and mutant. Repair-repressed: genes that are robustly transcribed only in mutant

Correlation analysis of viral read counts between CHX treatment of n12 and repair revealed that CHX led to a relatively stronger increase of Pol activity on IE genes on n12, compared to repair (Fig. 3C). DeSeq2 (17) fold change analysis also revealed Pol activity on IE and pre-IE genes, especially L/ST, were only minimally affected by CHX treatment in repair (Fig. 3D).

In contrast, CHX treatment in n12 infected cells led to a stronger increase on IE genes relative to other classes (Fig. 3E). Furthermore, we noted that the E and L genes of n12 that had significant increases in Pol activity were in genomic regions close to or nested with IE genes; (e.g., US10/11 nested with US12, and UL55 immediately downstream of UL54).

These findings indicated that virion components in the repair virus retain the ability to repress much of the aberrant transcription that occurs in n12 infection. Previous reports have shown ICP4 to be a strong repressor of L/ST (18,19) and therefore suggest ICP4 as the strongest candidate responsible for this repression. This was visible in the PRO-Seq data tracks since CHX treatment of repair yielded no visible increase in activity on L/ST and only negligible increases on the IE genes, α0, α4, and US1 (Fig. 3F). In contrast, CHX treatment of n12 led to extensive increases of Pol activity across all IE genes indicating that n12 virion-associated proteins were insufficient to mediate full repression.

The PRO-Seq visualization also revealed strong PPP on IE genes in both n12 and repair (Fig. 3F). Spline interpolation analysis of the relative read density across genes indicated that in the presence of CHX, both n12 and repair had a significant reduction in PPP (Fig. 3G/H), suggesting that the increase in reads was at least partially due to increased release of Pol into the gene body. However, CHX did not alter PPP peaks on the repair-repressed genes of n12 (Fig. 3I). These data further suggest that proteins other than ICP4 must be synthesized to regulate early transcription, and particularly to regulate PPP and release to elongation on IE genes.

### Viral IE gene ICP22 is also involved in early Pol repression

The previously characterized role of the IE gene ICP22 in the down regulation of Pol processivity and PPP enhancement (13), suggested a possible role for ICP22 in early HSV-1 Pol repression. To investigate this possibility, we used a ΔICP22 mutant virus lacking the entire US1 coding region and its genetically restored repair, which were derived from the HSV-1(F) strain (20,21). HEp-2 cells were infected with the mutant, repair, or wild type HSV-1(F) virus and PRO-Seq performed on nuclei harvested at 1.5 hpi.

ΔICP22 bore increased Pol activity across its genome compared to its genetically repaired counterpart at 1.5 hpi (Fig. 4A). We note that ΔICP22 repair had a significant increase in Pol activity relative to its parent wild type strain HSV-1(F) virus. This may be attributed to the repair expressing relatively lower levels of IE proteins than WT (20), or to the secondary mutations in the BAC background – notably the deletion of oriL (22). Nevertheless, ICP22 clearly played an important role in repression inasmuch as activity was significantly lower in the ΔICP22 repair virus than in the congenic ΔICP22 mutant (P < 0.0001) (Fig. 4B). The overall difference in reads was not as great as between n12 (ICP4-) and its repair (Fig. S5) but the pattern of Pol activity across the ΔICP22 genome as viewed by IGV was strikingly similar to n12, and covered the entire genome on both strands – a pattern distinct from repair and WT.

**Figure 4:**
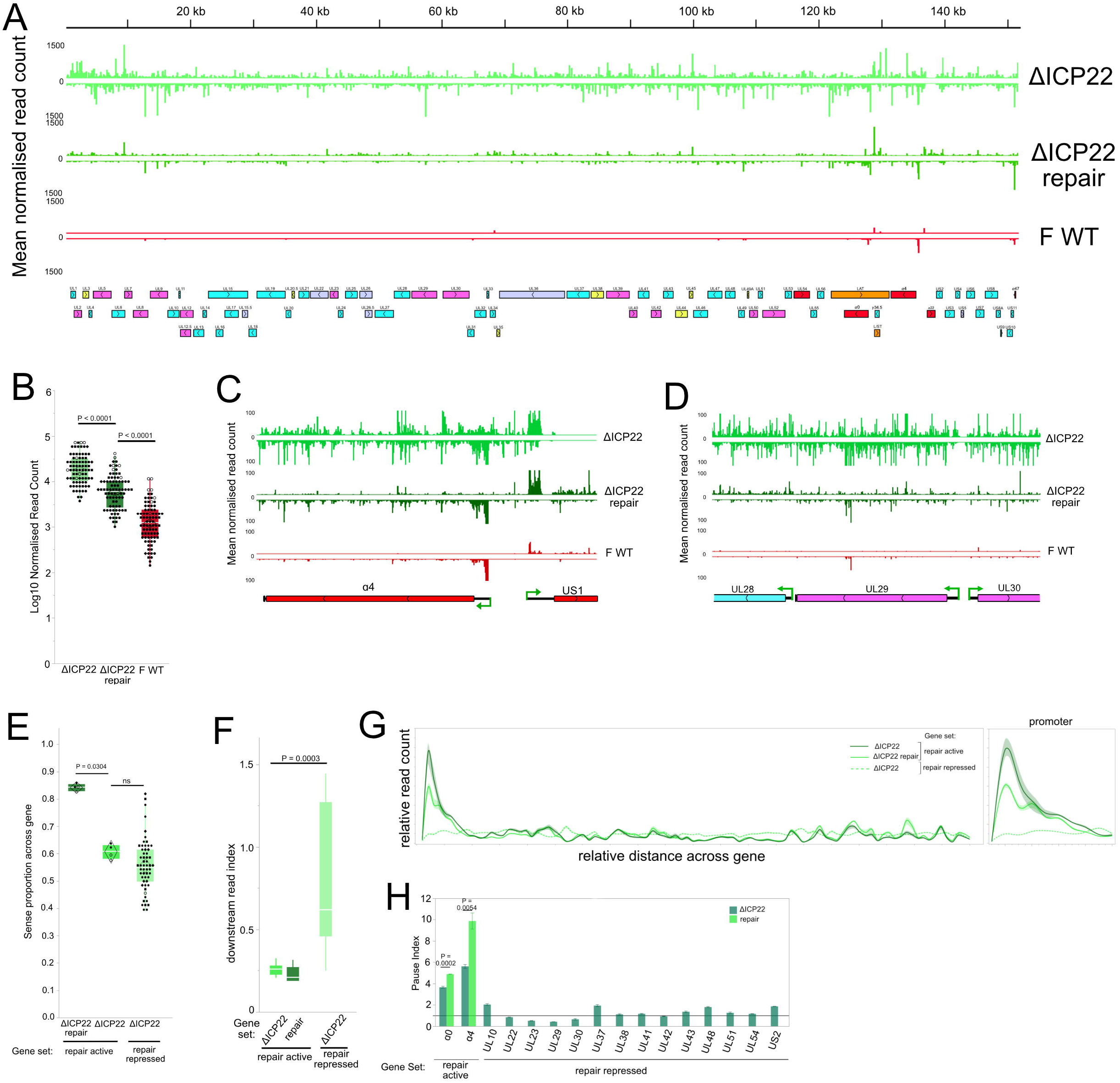
Aberrant viral transcription at 1.5 hpi in the absence of ICP22. HEp-2 cells were infected with a ΔICP22 mutant, its genetically restored repair and WT (F) and PRO-Seq performed at 1.5 hpi. **(A)** Genome browser view of distribution of PRO-Seq reads (data is mean of 3 biological replicates normalized to drosophila spike-in) across the HSV-1 genome. External repeat sequences were deleted for the sequencing alignment. **(B)** Log10 normalized read counts of all HSV-1 genes. High resolution genome browser view PRO-Seq tracks HSV-1 regions: **(C)** α4 and US1 and **(D)** UL28, UL29, UL30. IE genes: red, E genes: pink, LL genes: blue. **(E)** Proportion of reads mapping to the sense strand on repair-active genes. **(F)** Downstream-read index of repair-active and repair-repressed genes. Mann-Whitney U test was used to estimate statistical significance between mutant/repair or gene sets. ns = P > 0.05. **(G)** Spline interpolation analysis of the relative distribution of reads across repair-active and -repressed genes. A closer view of the promoter region is also shown. The bootstrap confidence of fit is shown in the shaded area. **(H)** Pause index values of repair-active and repair-repressed genes. Data is mean ± standard error. An unpaired student t-test was used to estimate statistical significance between mutant and repair on each gene. Repair-active: genes that are robustly transcribed in both repair and mutant. Repair-repressed: genes that are robustly transcribed only in mutant

Examples of dysregulated transcription were visible throughout the ΔICP22 genome. For example, Pol activity was increased at the α4/US1 region (Fig. 4C) and over the E genes UL29 and UL30 (which are not normally active at 1.5 hpi) (Fig. 4D). PRO-Seq patterns at these regions were inconsistent with properly regulated transcription with extensive antisense and intergenic transcription on the ΔICP22 genome and an apparent lack of PPP. The increase in antisense activity at 1.5 hpi was significant across all genes (Fig. 4E), indicating ΔICP22 lacked regulation of sense-to-antisense transcription across the entire genome.

Downstream-read index calculations also indicated a deficiency in transcriptional termination of the ΔICP22 virus, with significantly increased levels of reads downstream of the TTS on the repair-repressed genes (Fig. 4F). ΔICP22 also had a significant reduction in the promoter proximal peak on repair-active genes and an almost complete lack of this peak on repair-repressed genes at 1.5 hpi (Fig. 4G). This reduced PPP was confirmed by pause-index analyses (Fig. 4H). The lack of PPP on IE genes was reminiscent of the ΔICP22 phenotype noted previously at 3 hpi (13).

We previously showed that virion-associated ICP4 is required for full early transcriptional repression (Fig. 3). However, ICP22 is not known to be virion-associated (7) suggesting that ICP22 must be synthesized rapidly *de novo* to help suppress viral transcription. In order to investigate this possibility, we repeated the PRO-Seq experiment in the presence or absence of CHX. A small but significant increase occurred in reads across viral genes on both ΔICP22 and repair genomes after CHX treatment (Fig. 5A). However, the level of activity on genes classified as being robustly transcribed (read per bp ≥ 1 standard deviation of mean) did not change after treatment, indicating that the overall distribution of Pol across the viral genome was unchanged. This can be seen in the IGV visualisation in (Fig. 5B) with the repair retaining most of its Pol activity on IE genes. Further indication that the CHX treatment of the repair did not phenocopy the unrepressed ΔICP22 phenotype was indicated by the lack of effect on the proportion of sense transcription on the repair-active genes and a failure to lower sense transcription on the repair genome to the low level seen across the ΔICP22 mutant genome (Fig. 5C).

**Figure 5:**
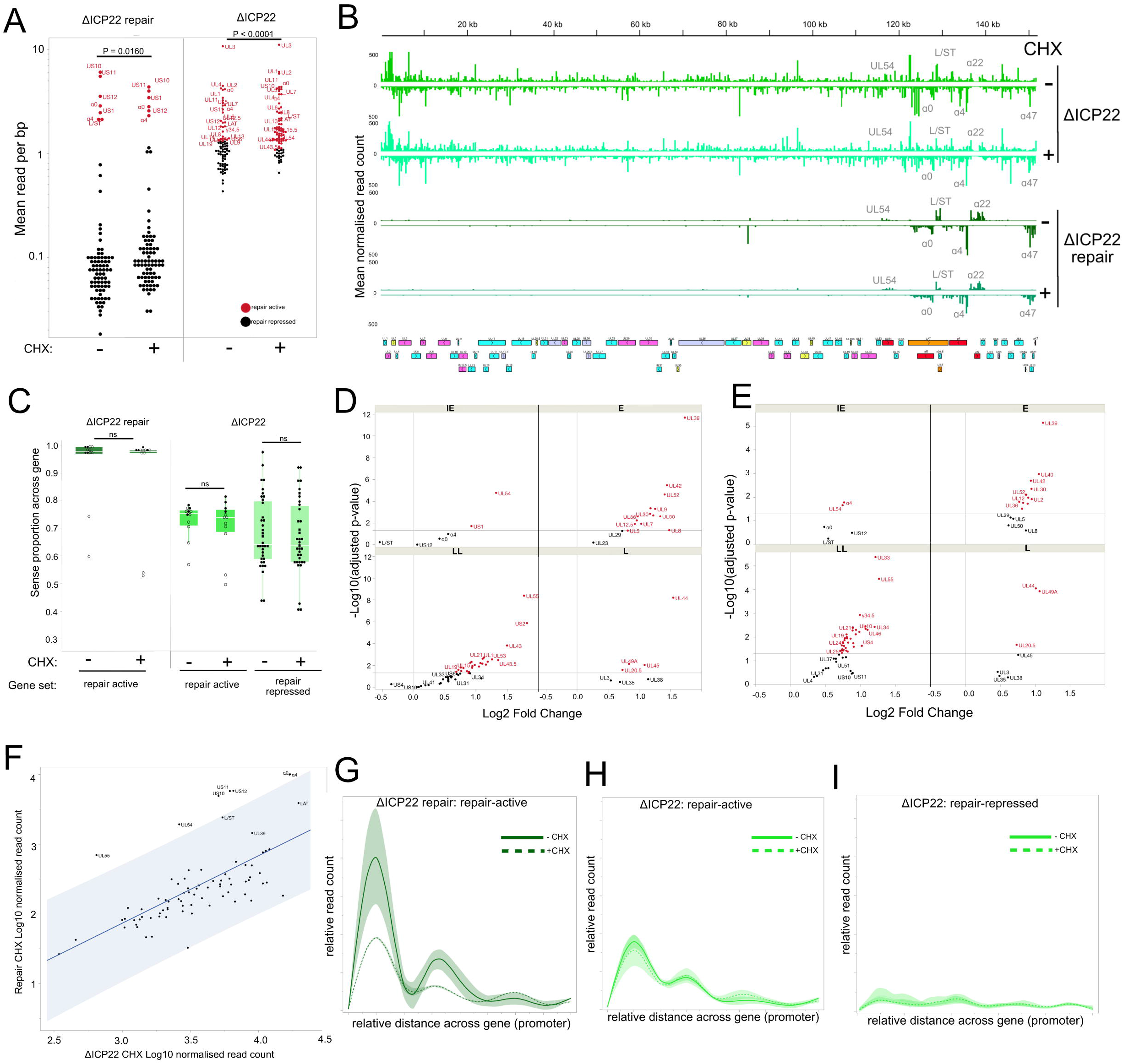
ICP22 is synthesized by 1.5 hpi to regulate PPP on IE genes. HEp-2 cells were infected with ΔICP22 mutant or its genetically restored repair in the presence or absence of the protein synthesis inhibitor cycloheximide (CHX) and PRO-Seq performed at 1.5 hpi. n=2 for each treatment, normalized based on total library read counts. **(A)** Mean read per bp across HSV-1 genes, red genes indicate those that were defined as being robustly transcribed **(B)** Genome browser view of distribution of PRO-Seq reads across HSV-1 genome, IE genes are indicated. **(C)** Proportion of reads mapping to the sense strand on repair-and repair-repressed genes. DeSeq2 Log2 fold change comparison of HSV-1 gene reads, separated by gene class of **(D)** repair CHX treated relative to repair untreated, **(E) Δ** ICP22 CHX treated relative to ΔICP22 untreated. IE: immediate early, E: early, LL: leaky late, L: late. Genes with a fold change adjusted p-value ≤ 0.05 are shown in red. **(F)** Scatterplot of HSV-1 gene reads after CHX treatment in repair and ΔICP22 infection. Shaded area indicates confidence prediction (mean ±standard error). Spline interpolation analysis of the relative distribution of reads across promoter regions in repair repair-active genes **(G)**, ΔICP22 repair-active genes **(H)** and ΔICP22 repair repressed **(I)**. The bootstrap confidence of fit is shown in the shaded area. Repair-active: genes that are robustly transcribed in both repair and mutant. Repair-repressed: genes that are robustly transcribed only in mutant

DeSeq2 Fold change analysis indicated that the read increases were not specific to a particular gene or class (Fig. 5D/E). Consistent with CHX treatment of n12 repair (Fig. 3D), IE genes were only minimally affected in the ΔICP22 repair, supporting the suggestion that virion components primarily repress transcription of these genes. Interestingly, correlation analysis of viral read counts between CHX treatment of ΔICP22 and repair revealed that CHX led to a stronger increase on IE genes (or those nested with IE genes) on repair relative to ΔICP22 (Fig. 5F), indicating virion components in ΔICP22 are more effective at repressing IE gene transcription. CHX treatment led to reduced PPP on repair-active genes on the repair (Fig. 5G) but did not alter the already reduced PPP on either repair-active or repair-repressed ΔICP22 genes (Fig. 5H/I). Overall, these data suggest that ICP22 is required to be rapidly synthesized by 1.5 hpi in order to regulate PPP on IE genes at early time points.

### Viral IE gene ICP0 is also required for early Pol repression

The CHX experiments have indicated that incoming proteins in wild type virions are sufficient to repress much of the aberrant transcription that occurs early in infection with IE mutants. Aside from ICP4, ICP0, is the only other known IE gene present in the virion (7). Therefore, we performed PRO-Seq in HEp-2 cells infected with HSV-1 n212, which contains a nonsense mutation in α0 derived from the KOS HSV-1 strain (23). To ensure the ICP0 control virus had a genetically identical background to the mutant, a repair virus was generated by recombination with α0 DNA. Immunoblotting was used to confirm repair of the ICP0 locus (Fig. S1D), and single-step growth curve analysis showed comparable replication kinetics and final yield of the repair relative to WT KOS (Fig. S1E/F). HEp-2 cells were infected with n212 or repair and PRO-Seq performed on nuclei harvested at 1.5 hpi.

As with n12 and ΔICP22, infection with n212 led to a statistically significant (P < 0.0001) increase in viral reads, relative to repair (Fig. 6A). The most striking difference in PRO-Seq profiles between n212 and its repair was an increase in activity on repair repressed genes at this early time point. For example, the E genes UL29 and UL30 are not normally active at

**Figure 6:**
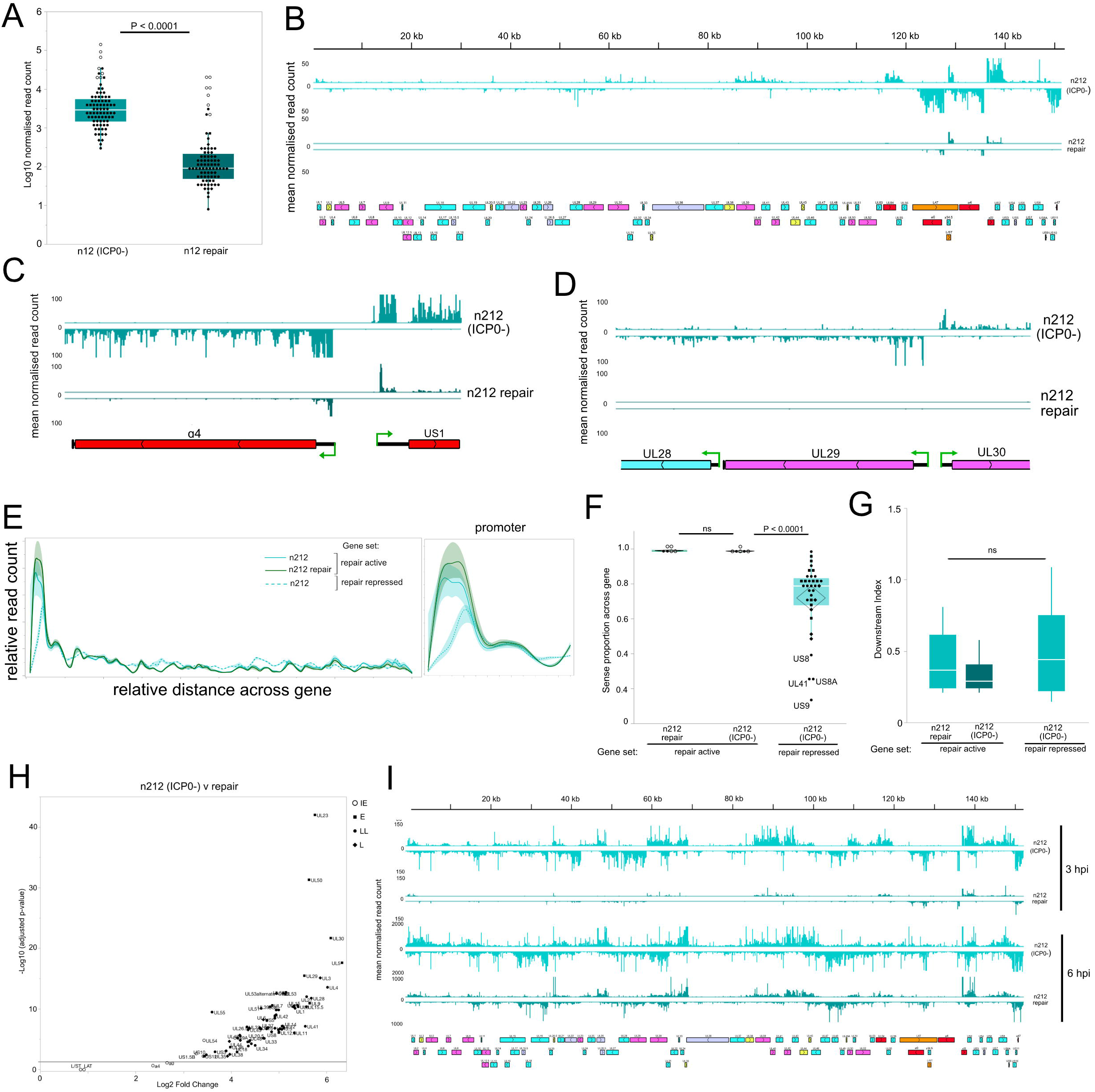
The absence of ICP0 leads to increased transcriptional activity on the HSV-1 genome at 1.5 hpi. **(A)** Log10 normalized PRO-Seq read counts of HSV-1 genes at 1.5 hpi post infection with n212 ICP0-mutant and its genetically restored repair (n=2, normalized to drosophila spike-in). Mann-Whitney U test was used to estimate statistical significance between mutant/repair. **(B)** Genome browser view of distribution of PRO-Seq reads across HSV-1 genome at 1.5 hpi. High resolution genome browser view PRO-Seq tracks HSV-1 regions: **(C)** α4 and US1 and **(D)** UL28, UL29, UL30. IE genes: red, E genes: pink, LL genes: blue. **(E)** Spline interpolation analysis of the relative distribution of reads across repair-active and -repressed genes. A closer view of the promoter region is also shown. The bootstrap confidence of fit is shown in the shaded area **(F)** Proportion of reads mapping to the sense strand on repair-active and repair-repressed genes. **(G)** Downstream-read index of repair-active and repair-repressed genes. **(H)** DeSeq2 Log2 fold change comparison of HSV-1 gene reads at 1.5 hpi of ICP0-mutant n212 infected relative to repair infected. **(I)** Genome browser view of distribution of PRO-Seq reads across HSV-1 genome at 3 and 6 hpi. Mann-Whitney U test was used to estimate statistical significance between viruses or gene sets at each time point. ns = P > 0.05. Repair-active: genes that are robustly transcribed in both repair and mutant. Repair-repressed: genes that are robustly transcribed only in mutant.

1.5hpi (as visible in repair) but bore substantial Pol activity on the n212 genome. However, the pattern of transcriptional activity more closely approximated that of normal regulation, with activity limited to ORFs and not covering the entire genome (Fig. 6B) as was seen in n12 and ΔICP22. For example, the PRO-Seq pattern of IE genes α4 and US1 included prominent PPP peaks and minimal antisense/intergenic activity (Fig. 6C). Even genes such as UL29 and UL30 that were aberrantly active in n212 displayed features indicative of proper transcriptional reregulation (Fig. 6D). Spline interpolation analysis confirmed that promoter peaks on repair-active genes in n212 were unchanged relative to repair, and that repair-repressed genes also maintained a PPP peak (Fig. 5E).

Quantification of sense/antisense transcription revealed that n212 had almost identical levels of sense transcription to repair, on repair-active genes (Fig. 6F). There was a reduced proportion of sense transcription on the repair-repressed gene set, but the proportion of sense transcription was higher than either n12 or ΔICP22. In addition, antisense transcription occurred heavily on US8, US8A, US9 and UL41, which bore high levels of antisense transcription in n12 and its repair (at 3 and 6 hpi) (Fig. 2F), further indicating that the antisense activity on these genes is a consistent feature of active HSV-1 transcription. In contrast to n12 and ΔICP22, downstream-read index calculations indicated that n212 retained normal transcriptional termination even on repair-repressed genes (Fig. 6G).

DeSeq2 analysis of individual genes indicated that n212 displayed the strongest increases in activity on specific E genes (Fig. 6H). There was no apparent trend for upregulation of a specific gene class over others in either n12 or ΔICP22 (Appendix 2). One interpretation of these data is that in the absence of ICP0, transcription progresses more rapidly through the temporal transcriptional program, causing later genes to be transcribed earlier. PRO-Seq data using n212 at 3 and 6 hpi was consistent with this hypothesis, inasmuch as the distribution of Pol across the genome in n212 at 1.5 hpi matched that of the repair at 3 hpi (Fig. 6I). By 6 hpi, n212 and its repair had virtually matching PRO-Seq profiles, although n212 continued to have higher read counts. This result differed from ΔICP22 which had reduced Pol activity on IE genes at 3 hpi and loss of Pol activity from most genes by 6 hpi (13), and n12 which primarily retained activity on only IE genes at 3 and 6 hpi (Fig. 1).

To determine whether the increased PRO-Seq reads during infection with IE gene mutants was a result of increased genome copies entering the nucleus, we used qPCR to quantify viral DNA levels in isolated nuclei at 1.5 hpi. All mutants did have increased numbers of incoming genomes relative to their congenic repairs (Fig. S5). However, the increased genome copies could not fully account for the increase in PRO-Seq reads aligning to the mutant viral genomes. Therefore, much of the increased transcriptional activity of the mutants was attributable to repression mediated by the respective IE genes rather than changes in particle infectivity.

## Discussion

We have shown that ICP0, ICP4 and ICP22 are individually required for early transcriptional repression on the HSV-1 genome that precedes the activation of the temporal transcriptional cascade. High Pol activity might be expected early in infection because HSV-1 genomes enter the nucleus free of nucleosomes and contain densely packed, mostly intronless genes encoded on both strands. This gene arrangement necessitates a high concentration of promoter elements such as TATA boxes and initiator sequences recognized by cellular Pol (24,25), and multiple GC-rich regions that can bind cellular transcription factors including Sp1 and Egr-1 (26,27). In addition, VP16 co-introduced with the genome is a powerful transcription factor with potent DNA binding activity for viral IE promoters, and a transcriptional activation domain (TAD) that interacts with many transcription factors such as TFIIA, TFIIB, TBP and TFIIH (28). The effectiveness of these elements is supported by the observation that by 3 hours after infection, the viral genome bears 1/3 of all Pol activity of the cell (29).

It is likely that the newly introduced HSV-1 genome is rapidly engaged by not only virion-delivered VP16, ICP0 and ICP4, but also by cellular histones. Previous studies have shown that histones associate with the viral genome during lytic infection (reviewed in (30)) and that histone dynamics are important in the regulation of lytic transcription (31). Other studies emphasize the ability of ICP4 to coat the viral genome in a form of “viral chromatin” (32).

ICP4 in this viral chromatin was proposed to facilitate recruitment of Pol and components of the pre-initiation complex. However, because Pol is efficiently recruited to the viral genome in the absence of full-length ICP4 (and ICP0 and ICP22) we propose that a major function of ICP4 and virally-orchestrated chromatin is to transiently repress transcription.

The repressive role of ICP4 is consistent with ICP4’s known ability to repress IE gene transcription and was supported by our 6 hpi PRO-Seq data, in which n12 was unable to repress IE gene Pol activity. ICP4 repression is associated with ICP4-binding sites (18,19,33), that have a relatively loose consensus sequence with more than 100 copies throughout the viral genome (32). CHX treatment showed that virion-associated ICP4 represses IE genes preferentially. This may seem paradoxical since the tegument also contains the major IE transcriptional activator, VP16 (2), but further highlights the intricate balance of control the virus exerts over its own transcription.

It is possible that early transcriptional repression is dependent on the form of ICP4, which must switch from a repressor to an activator on certain genes at certain times. n12 is not completely null for ICP4 and expresses a truncated form (Fig. S1A), containing most of the N-terminal activation domain but lacking the DNA-binding domain (35). The N-terminal domain has been shown to be required for viral gene activation through interactions with TFIID and mediator (36). Therefore, the abundant transcription in n12 at 1.5 hpi may be due to this truncated form of ICP4 recruiting transcription factors and its inability to form a complex with TFIIB and TBP, which is believed to foster repression (37).

The result with ΔICP22 likely reflects ICP22’s known role in reducing processivity of RNA Polymerase (13), yet it was surprising that this occurred over the entire viral genome and was not limited to IE genes. One possible explanation is that transcription is initiated at VP16 sites, but that the increased Pol processivity in the absence of ICP22 leads to substantial read through into neighbouring genes (38). ΔICP22 mutant virions have a defect in virion composition (34), including an increase in ICP4, and could also therefore contain an increased level of VP16. This increase in ICP4 may also explain why the ΔICP22 mutant was more effective at IE repression than its repair (Fig. 5F) and support the finding that tegument ICP4 primarily functions to repress IE gene transcription. ICP22 has also been shown to interact with several elongation regulatory factors including FACT (39), p-TEFb (40) and CDK9 (41) and it has been proposed that these interactions leads to selective repression of cellular genes (42). A similar mechanism might account for repression of the HSV-1 genome early in infection.

We also noted an increase in early transcription in cells infected with the α0 mutant n212. ICP0 is enriched in ICP22 mutant virions (34) and has an increased association with n12 genomes, relative to WT (16). While ICP0 functions as a promiscuous transactivator that acts by causing degradation of cellular transcriptional silencing factors (43) PRO-Seq of cells infected with n212 unexpectedly exhibited increased transcriptional activity at 1.5 hpi.

Unlike the ICP4 and ICP22 mutants, the pattern of Pol activity on the ICP0 mutant genome reflected normal regulatory features. It is known that ICP0 mutants infected at high MOI grow equivalently to wild-type virus (43)(can be seen in Fig. S1F) and that these mutants produce a large number of defective particles at very early stages of infection (23). One possibility is that the large number of defective particles deliver more abundant virion components such as the incoming VP16/ICP4 to help activate the n212 genome. Our CHX experiments also highlighted the importance of virion components in early transcriptional repression

HSV-1 forms membrane-less liquid-like transcriptional condensates (44) and ICP4 has been identified as an important factor for their formation (45). It is possible that these condensates act to protect the viral genome from unregulated transcription. Promyelocytic leukemia nuclear bodies (PML NBs or ND10) are also liquid-like membrane-less organelles. ND10 components rapidly associate with incoming HSV-1 genomes, presumably to repress infection (46), and are usually targeted for proteasomal degradation by ICP0. Disruption of IE gene expression may alter the formation of these condensates, impairing recruitment of repressors to the viral genome and leading to unfettered exposure to the cellular transcriptional machinery resulting in rampant Pol activity.

The lack of transcriptional regulation across most of the n12 and ΔICP22 genomes at 1.5 hpi suggests that co-transcriptional processing events are also disrupted (38). Consequently, it is likely that the RNA generated is not processed into mature mRNA, though this remains to be determined. The large amount of RNA being produced is a potential trigger for anti-viral responses and indicates why rampant transcription is likely disadvantageous to a virus such as HSV-1. This is supported by the finding that both n12 and ΔICP22 act to repress this aberrant transcription by 3 and 6 hpi. We propose that the timing of this repression may explain some of the differences between experiments. For example, the overall read difference between n12 and repair was less extensive when repeating the experiment with CHX and in HFF cells, suggesting the virus had begun to gain control of transcription already. The fact that the unrepressed phenotype was strongest at just 1.5 hpi likely explains why this process has not been previously identified. Furthermore, we were only able to identify the requirement for early transcriptional repression by comparison to ICP4 and ICP22 mutants, which other HSV-1 nascent RNA-Seq studies did not include (47). We also note that PRO-Seq identifies nascent transcription that does not necessarily correlate with mature mRNA accumulation. Therefore, mRNA-Seq studies (6) would not necessarily identify the increased Pol activity shown here.

Because each IE protein affects production of the others, we cannot currently determine whether the observed de-repression was due to the specific loss of one, or a combinatorial effect, and the full mechanism of repression remains to be elucidated. Nevertheless, we have shown that prior to initiating activation of transcription, IE genes first mediate transcriptional repression. This is a previously unrecognized process preceding steps outlined in the current paradigm of HSV transcription which proposes that IE gene products function primarily to promote viral gene expression (48). These data suggest that the current paradigm be modified to consider the entire cascade of viral gene expression as a series of de-repression steps on different genes. This transient repression may represent an important checkpoint for a virus that can orchestrate one of two potential transcriptional programs: one that is robust during productive replication in which repression is reversed, or one during latency in which repression is maintained.

## Materials and Methods

### Cells

HEp-2 (human epithelial lung cancer), Vero (African green monkey), RSC (rabbit skin) and the Vero derived ICP4-complementing, E5, cells (49) were maintained in Dulbecco’s modified Eagle’s medium (DMEM) containing 10% new born calf serum (NBS), 100 units/ml penicillin, 100μg/ml streptomycin (pen/strep) and maintained at 37 □ C with 5% CO_2_. U2OS (human bone osteosarcoma) cells were maintained in McCoy’s 5A medium containing 10% foetal bovine serum (FBS), pen/strep and maintained at 37 □ C with 5% CO_2_. HFF (human foreskin fibroblasts) were maintained in DMEM containing 15% FBS, pen/strep and at 37 □ C with 5% CO_2_. S2 (*Drosophila melanogaster*) cells were grown in Schneider’s medium containing 10% FBS and maintained at 23°C. E5 cells were a gift from Dr. Neal DeLuca, University of Pittsburgh and HFF cells were a gift from Dr. Luis Schang, Cornell University.

### Viruses

Mutant HSV-1 strains n12 (49) and n212 (23), both HSV-1 KOS derived, were gifts from Dr. Neal DeLuca, University of Pittsburgh. n12 virus stocks were prepared on E5 cells and n212 were prepared on U2OS cells. The ΔICP22 virus and its repair, generated from a HSV-1 (F) bacterial artificial chromosome, were gifts from Dr. Yasushi Kawaguchi (20,21). Stocks were prepared on Vero cells.

Repair viruses of n12 (α4 mutant) and n212 (α0 mutant) were generated through homologous recombination. Plasmids containing WT HSV-1 DNA spanning 1000 bp either side of the mutation were synthesized, linearized, then transfected using TransIT-X2 (Mirus Bio) into RSC cells. 6 hours after transfection, cells were infected at a MOI of 1 with the corresponding mutant and when all cells showed CPE, harvested by scraping into media.

Viruses were released from cells through 3x cycles of freeze-thaw in LN_2_ and 37°C water bath. Samples were then serially diluted onto Vero cells with an agarose overlay and underwent 3 rounds of plaque purification. Repair virus plaques were selected by size due to the growth restriction of the mutants on Vero cells. Western blotting was used to confirm successful repair of full-length protein expression (Fig. S1). Stocks were subsequently prepared on Vero cells.

### Virus infections and drug treatments

For assessment of IE protein expression, monolayers of monolayers of HEp-2 cells were infected with HSV-1 at a MOI of 5 in 199V medium (+1% NBS). After 1h, inoculum was replaced with DMEM (+2% NBS).

For PRO-Seq, monolayers of HEp-2 cells were infected with HSV-1 at a MOI of 5 as above. Infections were allowed to continue for 1.5, 3 or 6 hpi – hpi refers to the time after the viral inoculum was first added to cells. For CHX treatments, 10μm of CHX was added to media 1h prior to infection and this CHX concentration was maintained in the media throughout infection.

### Single-step growth curve analysis

Monolayers of HEp-2 cells were infected with HSV-1 at an MOI of 5 in 199V medium (+1%NBS) at 37°C. After 1h, inoculum was removed, cells were washed 3x in PBS to remove unabsorbed virus and DMEM (2% NBS) was added. Infected cells were harvested at 0, 6, 12, 18 and 24 hpi by scraping into media and viruses released from cells by 3x cycles of freeze-thaw.in LN_2_ and 37°C water bath. Virus yields were determined by plaque assay on Vero cells or appropriate complementing cell-line.

### Western blotting analysis

Lysates were prepared from infected cells by washing cells x2 in ice-cold PBS before addition of an appropriate volume of NP-40 lysis buffer (150mM NaCl, 1% NP-40, 50mM Tris pH8, protease inhibitors). Cells were then scraped into suspension and incubated on ice for 30 min. 20µg of protein was mixed 2X Protein Sample Loading Buffer (LiCor) and heated to 98 □ C for 5 min. Samples were separated on an 8% polyacrylamide resolving gel, layered with a 5% stacking gel. Proteins were transferred to a 0.45 μm nitrocellulose membrane using a wet transfer technique. Membranes were first blocked for 1 h at room temperature in 5% BSA +0.1% Tween _®_ 20, then incubated with the primary antibodies diluted in blocking buffer at 4 □ C overnight. Membranes were washed 4 x for 5 min in PBS containing 0.1% Tween before being incubated with the secondary antibodies diluted in blocking buffer at room temperature for 45 min. Membranes underwent a further 4x 5 min washes in PBS-T and were then visualized on an Odyssey Scanner (LiCor). Primary antibodies were anti-ICP4 (sc-69809, Santa Cruz), used at 1:200 and anti-ICP0 (sc-53070, Santa Cruz), used at 1:100. Secondary antibodies were DyLight 680 and 800 (Cell Signalling).

### Nuclei isolation and PRO-Seq

Nuclei were isolated from infected cells, nuclear run-on with biotinylated nucleotides performed and sequencing libraries generated as described previously (5,13,29,50). Nuclei from Drosophila S2 cells were spiked into infected-cell nuclei at a ratio of 1:1000 prior to run-on. Libraries were sequenced on an Illumina NextSeq 500, performed by the GeneLab at Louisiana State University School of Veterinary Medicine or by the Biotechnology Resource center (BRC) Genomics facility *(RRID:SCR_021727)* at the Cornell Institute of Biotechnology.

### DNA isolation

The lower interphase and organic layers of TRIzol LS RNA extracted HEp-2 infected nuclei was saved. The DNA was subsequently isolated following the manufacturer’s protocol for TRIzol DNA isolation (Invitrogen, MAN0016385). To clean-up the isolated DNA, 3x phenol:chloroform:isoamyl alcohol purification was performed before ethanol precipitation using sodium acetate. DNA was resuspended in DEPC water.

### Quantitative PCR for viral genome copy

Extracted DNA was quantified using NanoDrop spectrophotometer (Thermo Scientific). HSV-1 genome copy number was determined in 50ng of extracted DNA by qPCR as previously described (5,29) using Brilliant III SYBR _®_ Green Master Mix with ROX (Agilent); and analysis was performed using QuantStudio^(tm)^ 3 (Applied Biosystems).

### Read processing

FastQ files were processed using the PRO-Seq pipeline developed by the Danko lab (Cornell) https://github.com/Danko-Lab/utils/tree/master/proseq. Reads were aligned to a concatenated genome file containing hg38, dm3, HSV-1 genomes. HSV-1 genome builds had the external repeats deleted to aid sequencing alignment; the modified genome files are available: https://github.com/Baines-Lab/Public/tree/main/HSV-1. Seqmonk software (51) was used to probe reads from the individual genomes in the output .bam files. Drosophila spike-in normalisation was used to account for variation in sequencing depth between libraries.

Libraries were normalized relative to the library with the largest drosophila read count, with the scale factor for this set to 1. Libraries prepared without spike-in were normalized based on total library read counts. The output .bw files (containing only the 3’ final read position) were visualized using the IGV genome browser (52). Individual reads for each bp across the HSV-1 genome was extracted from the .bw files using multiBigWigSummary from deepTools (53) and normalized as above. The resulting .txt files were used for analysis in Seqmonk or .bedgraph for visualization in IGV. The normalized .txt/.bedgraph files were used for all subsequent data analysis.

### Identification of transcriptionally active genes

The robust transcription threshold to identify repair-active genes was determined on data from repair-virus infection at 1.5 hpi due to restricted transcriptional activity on these samples. Genes were classified to be transcriptionally active (repair-active) if the mean read per bp (TSS -TTS) was greater than 1 standard deviation of the mean of all genes. Genes below this threshold in repair infections, but above this threshold in mutant infections were classified as repair-repressed. The same threshold was used at all time points and was calculated individually for each sequencing experiment due to variation in sequencing depth. Details of normalized HSV-1 gene reads, reads per bp and robust transcription thresholds are given in Appendix 1.

### Promoter proximal pause analysis

The relative distribution of reads across gene sets was determined using the Seqmonk probe trend plot on .txt files. Genes were divided into 100 bins to get relative distance across genes. Data was plotted using the smoother function in SAS JMP Pro software and the bootstrap confidence of fit calculated using 300 iterations of the data. Pause index calculations for individual genes was determined by the formula: mean read per bp (TSS +150bp)/mean read per bp rest of gene (to TTS). Only viral genes with no overlapping transcripts and with a defined TSS were included in PPP analysis to allow reads to be accurately assigned.

### Sense transcription proportion calculation

The proportion of sense transcription on each HSV-1 gene was calculated by formula: reads sense strand of gene (TSS-TTS)/reads antisense strand of gene (TSS-TTS). Only isolated HSV-1 genes without genes on the opposite strand were included in final analysis to allow reads to be accurately assigned. Regression analysis was used to assess correlation between sense proportion and read per bp (TSS-TTS) of each HSV-1 gene. The square of the correlation coefficient (R^2^) was calculated using SAS JMP Pro software.

### Downstream-read index calculation

The level of reads downstream of TTS was calculated by the formula: reads per bp 150bp downstream of TTS/reads per bp in upstream ORF (TSS-TTS). Only singular unnested genes with a defined TTS more than 150bp away from any other TSS were included to allow reads to be accurately assigned.

### Fold change analysis

Fold change analysis was performed using the R package DeSeq2 (17). DeSeq2 for HSV-1 genes was performed incorporating Hg38 genes data to account for the library size correction step. Full DeSeq2 fold change and p-values are given in Appendix 2.

### Statistical analysis

Statistical significance was determined by unpaired Student’s t-test for pairwise comparisons of normally distributed data, Mann-Whitney U test was used for pairwise comparison of nonparametric data and Kruskal-Wallis for multiple comparisons. Calculations were performed using SAS JMP Pro software. Each independent experiment consisted of 2-3 biological replicates.

## Data availability

The data will be publicly available upon publication on the GEO database under the accession numbers GSE202363 and GSE213098. Reviewers can access the data with secure tokens groxeaugddedhqr and ujcbyqmwnrgvfer, respectively.

## Acknowledgements

We thank Dr. Neal DeLuca, Dr. Luis Schang and Dr. Yasushi Kawaguchi for their kind gifts of cells and viruses. We thank Thaya Stoufflet and Dr. Vladimir Chouljenko at the LSU SVM GeneLab and the Genomics Facility (*RRID:SCR_021727*) of the Biotechnology Resource Center of Cornell Institute of Biotechnology for their help with sequencing experiments.

Portions of this research were conducted with the high-performance SuperMic supercomputer provided by Louisiana State University (http://www.hpc.lsu.edu) and we thank Dr Le Yan for his help in maintaining the PRO-Seq pipeline. These studies were supported by National Institutes of Health grants R01 AI 141968 and R21 AI 148926 to J.D.B.

## Author contributions

J.D.B conceived the study, L.E.M.D performed experiments involving n12 and n212 HSV-1 viruses. C.H.B and R.D performed experiments involving ΔICP22 HSV-1. L.E.M.D performed data analysis and made the figures. All authors discussed results L.E.M.D and J.D.B wrote and edited the manuscript.

## Supplementary Information

**Figure S1: Successful generation of n12 and n212 repair viruses. (A)** Western blot of mock, n12 (ICP4-), n12 repair and WT (F) infected HEp-2 lysates using anti-ICP4 antibody. Single-step growth curve analysis of WT (KOS), n12 and n12 repair from infected HEp-2 cells (MOI:5) titrated by plaque assay on Vero cells **(B)** and the ICP4-complementing cell line, E5 **(C)**. Data is mean of 3 biological replicates ± standard error. **(D)** Western blot of mock, n212 (ICP0-), n212 repair and WT (F) infected HEp-2 lysates using anti-ICP0 antibody. Single-step growth curve analysis of WT (KOS), n212 and n212 repair from infected HEp-2 cells (MOI:5) titrated by plaque assay on Vero cells **(E)** and U2OS cells **(F)**.

**Figure S2: Pause indices for individual HSV-1 genes after infection with IE mutants and their congenic repair viruses**. Pause index values calculated from PRO-Seq data of repair-active (genes that are robustly transcribed in both repair and mutant) and repair-repressed (genes that are robustly transcribed only in mutant) HSV-1 genes in **(A)** ICP4-mutant n12 and n12 repair infection at 1.5 hpi (n=2), **(B)** 3 hpi (n=2) and **(C)** 6 hpi (n=2). **(D)** n12 and repair at 1.5 hpi, in the presence or absence of CHX (n=2 for each treatment). **(E)** ΔICP22 and its repair at 1.5 hpi in the presence or absence of CHX (n=3 for each treatment). **(F) I**CP0-mutant, n212 and its repair at 1.5hpi (n=2). Data is mean ± standard error. An unpaired student t-test was used to estimate significance between mutant and repair on each gene and are noted where significant.

**Figure S3: Antisense transcription of specific genes at regions downstream of ICP4-independent promoters**. High resolution PRO-Seq tracks (mean of 2 biological replicates, normalized to drosophila spike-in) of regions containing ICP4-independent promoters (shown by green arrows) on the HSV-1 genome after infection with ICP4-mutant n12 and its repair at 1.5, 3 and 6 hpi. **(A)** The UL54 ICP4-independentt promoter and UL56 antisense transcription. **(B)** The US1 ICP4-independent promoter and US2 antisense transcription. **(C)** The US12 ICP4-independent promoter and US8/US8A/US9 antisense transcription. **(D)** The Ul39 ICP4-independent promoter and UL41 antisense transcription. **(E)** The α0 ICP4-independentt promoter and **(F)** The α4 ICP4-independent promoter. Dashed lines define regions of potential read-through transcription.

**Figure S4: Aberrant transcription on the HSV-1 genome at 1.5 hpi in the absence of ICP4 in HFF cells**. HFF cells were infected with either the ICP4-mutant, n12, or its genetically restored repair and PRO-Seq performed at 1.5 hpi. **(A)** Genome browser view of distribution of PRO-Seq reads (mean of 2 biological replicates, normalized to drosophila spike-in) across the HSV-1 genome. External repeat sequences were deleted for the sequencing alignment. IE gene peaks are noted. **(B)** Log10 normalized read counts of all HSV-1 genes and **(C)** of genes separated by temporal class. **(D)** Proportion of reads mapping to the sense strand on repair-active (genes that are robustly transcribed in both repair and n12) and repair-repressed (genes that are robustly transcribed only in n12). genes. **(E)** Downstream-read index of repair-active and repair-repressed genes. Mann-Whitney U test was used to estimate statistical significance between mutant/repair or gene sets. ns = P>0.05. **(F)** The relative distribution of reads across repair-active genes and repair-repressed genes A closer view of the promoter region is also shown. The bootstrap confidence of fit is shown in the shaded area. **(G)** Pause index values of repair-active and repair-repressed HSV-1 genes. Data is mean of 2 biological replicates ± SEM.

**Figure S5: HSV-1 genome copy number in nuclei at 1.5 hpi in HEp-2 and HFF infected cells**. The number of HSV-1 genomes in infected cell nuclei at 1.5 hpi were determined via UL51 plasmid standard-curve qPCR in HEp-2 and HFF infected cells. Reported data comprises a mean of 2 biological replicates ± standard error. An unpaired student t-test was used to estimate statistical significance, ns = P>0.05. The table below indicates fold differences in viral genome copies, compared to fold differences in total HSV-1 reads in PRO-Seq experiments. ┼ indicates that genome copy was determined in an experiment independent of PRO-Seq. * indicates that genome copy was determined on nuclei matched to a PRO-Seq experiment.

**Appendix 1:** Normalized HSV-1 gene reads, reads per bp and robust transcription thresholds

**Appendix 2:** DeSeq2 fold change and p-values

## Notes

### Competing Interest Statement

The authors have declared no competing interest.

### Summary of Updates

Revised to include an additional experiment - CHX treatment during infection with ICP22 mutant.

