## Supplementary figures and images for "Immediate early proteins of herpes simplex virus transiently repress viral transcription before subsequent activation"

### Figure S1

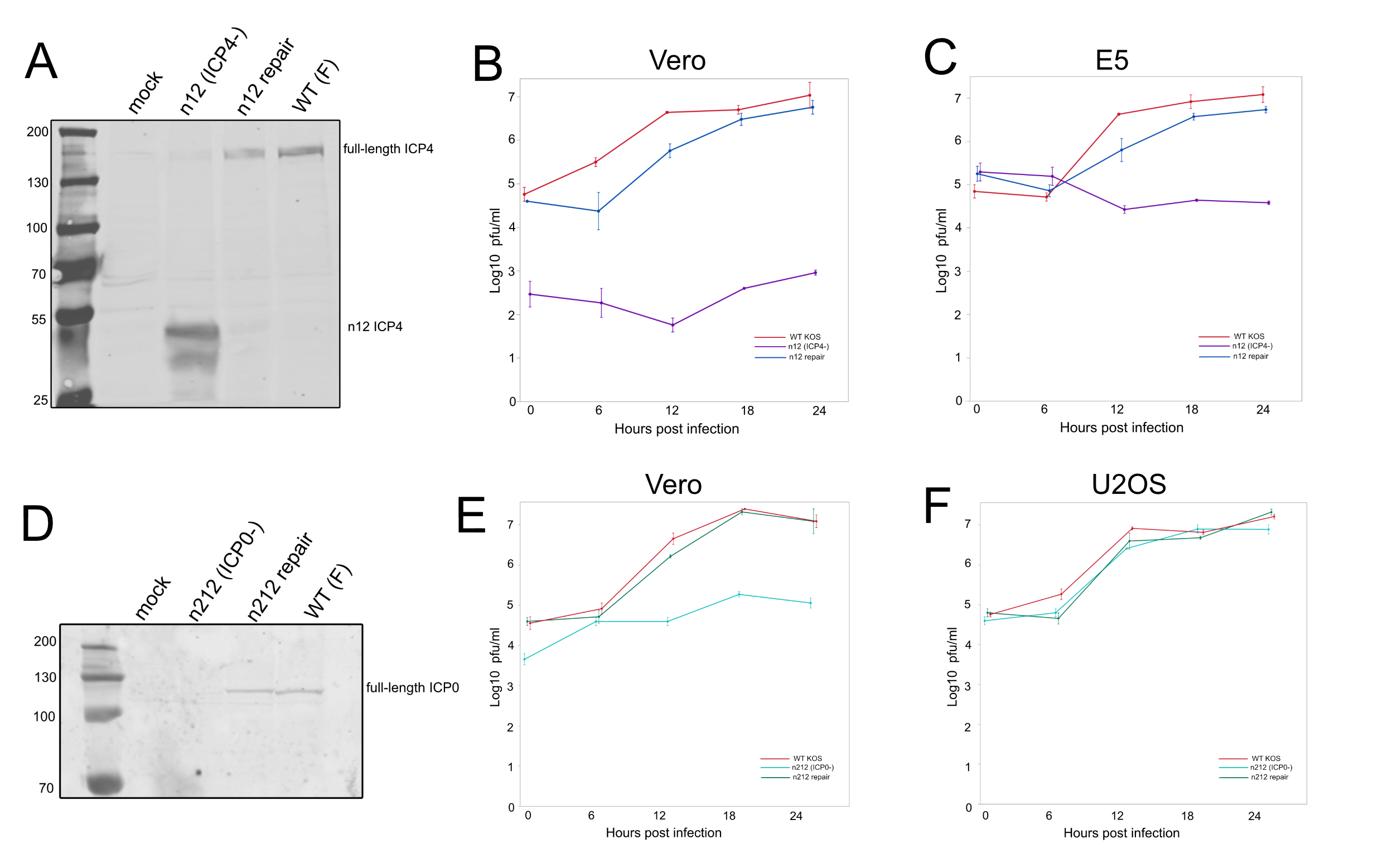

### Figure S2

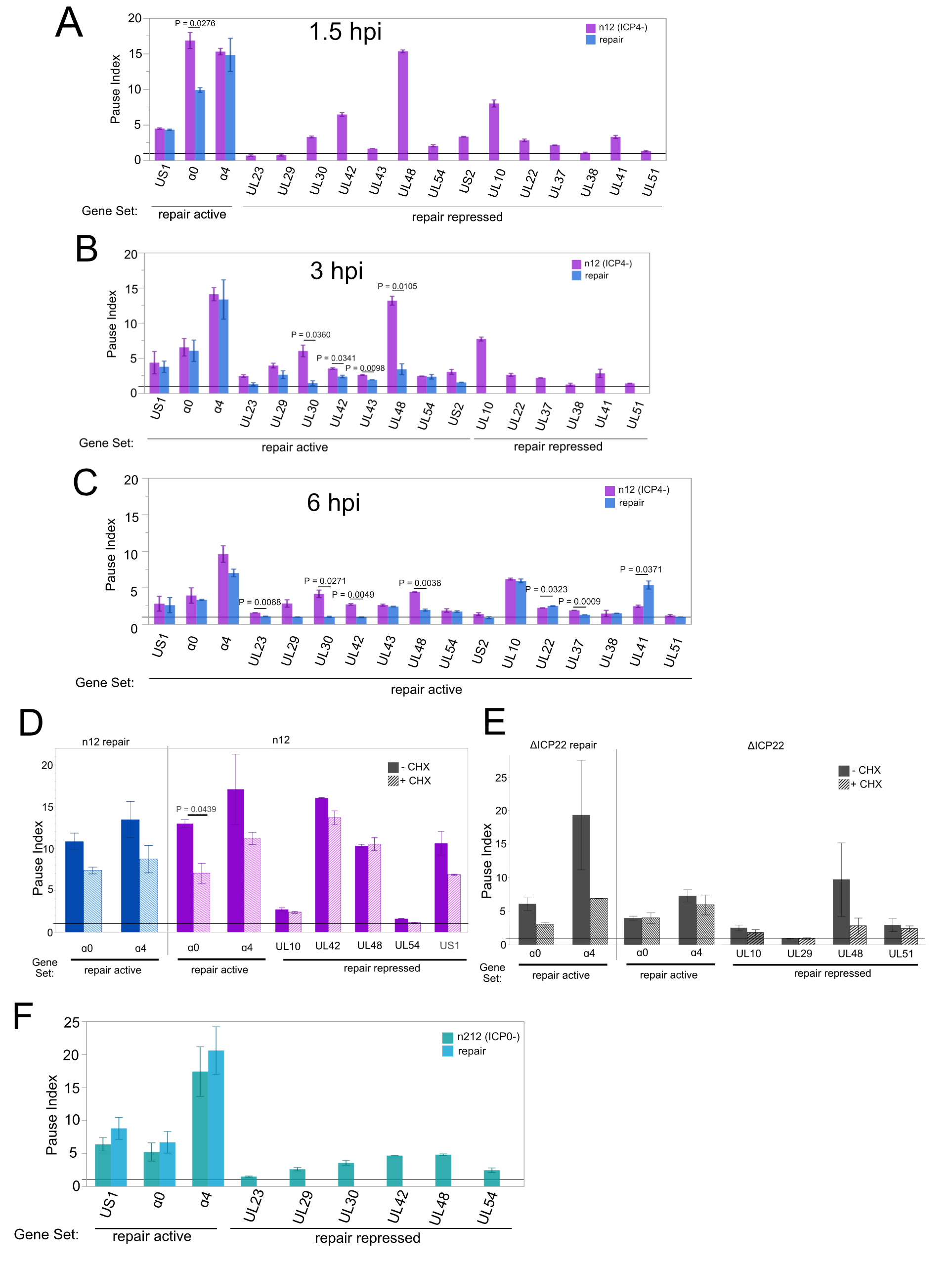

### Figure S3

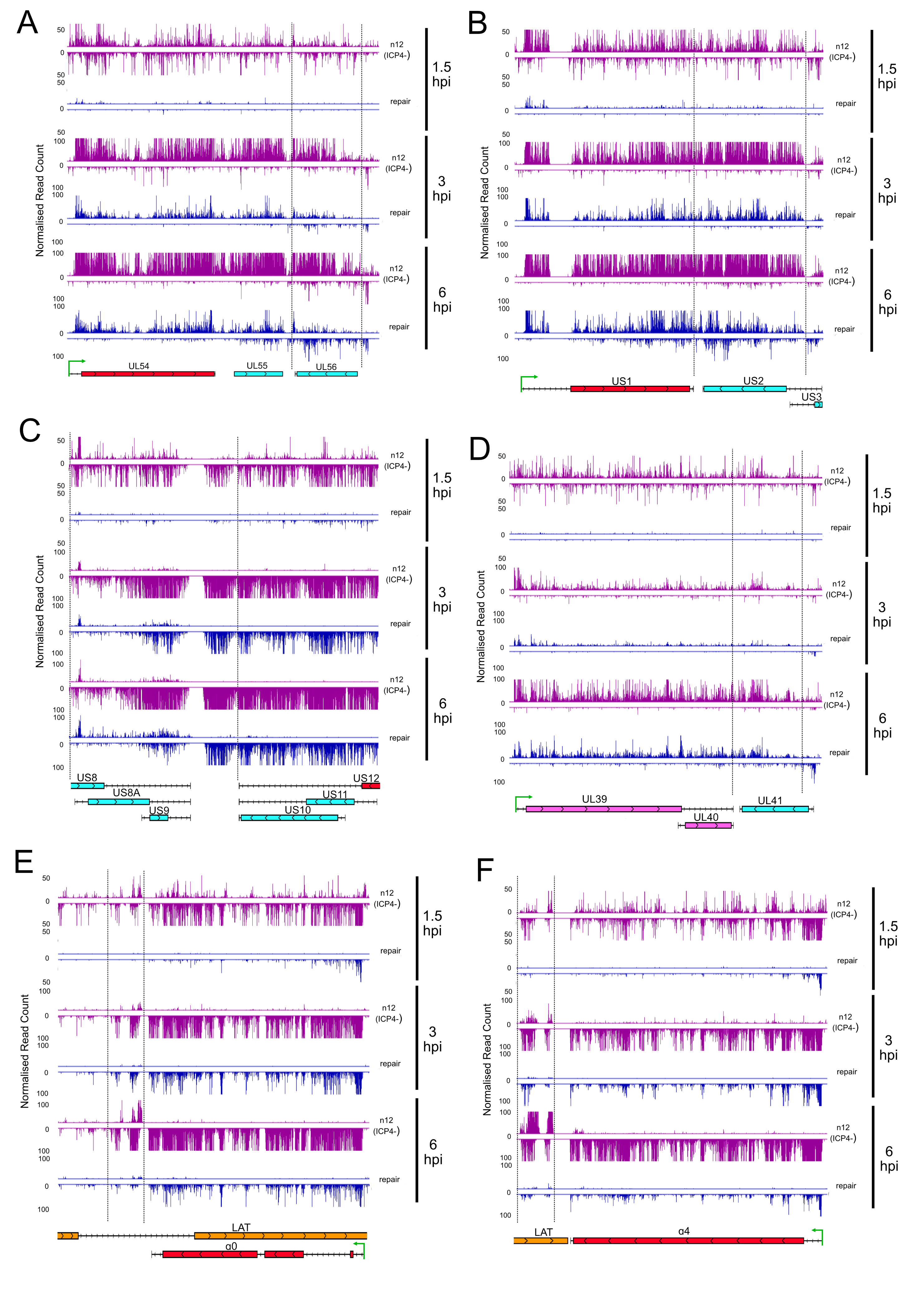

### Figure S4

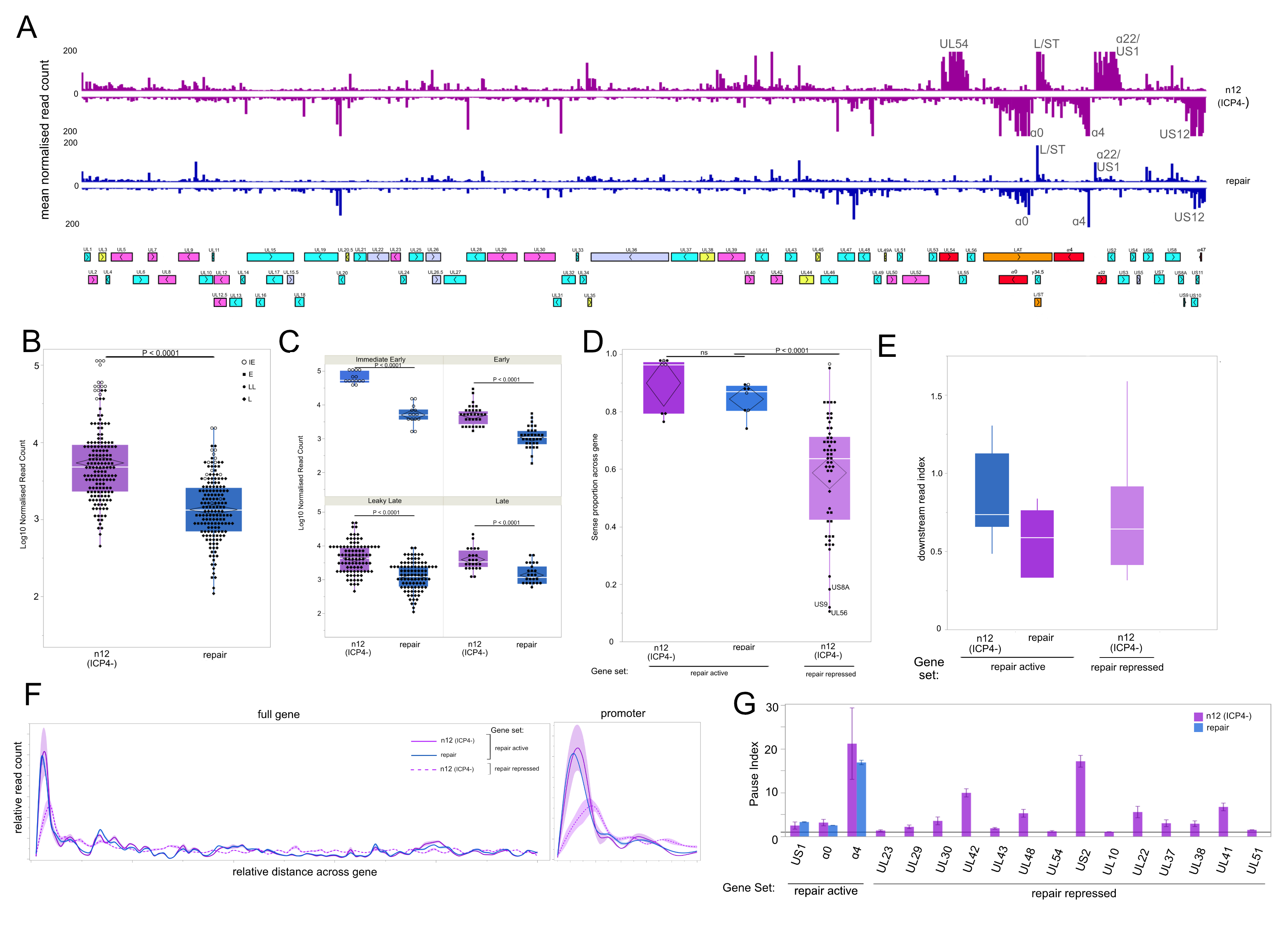

### Figure S5

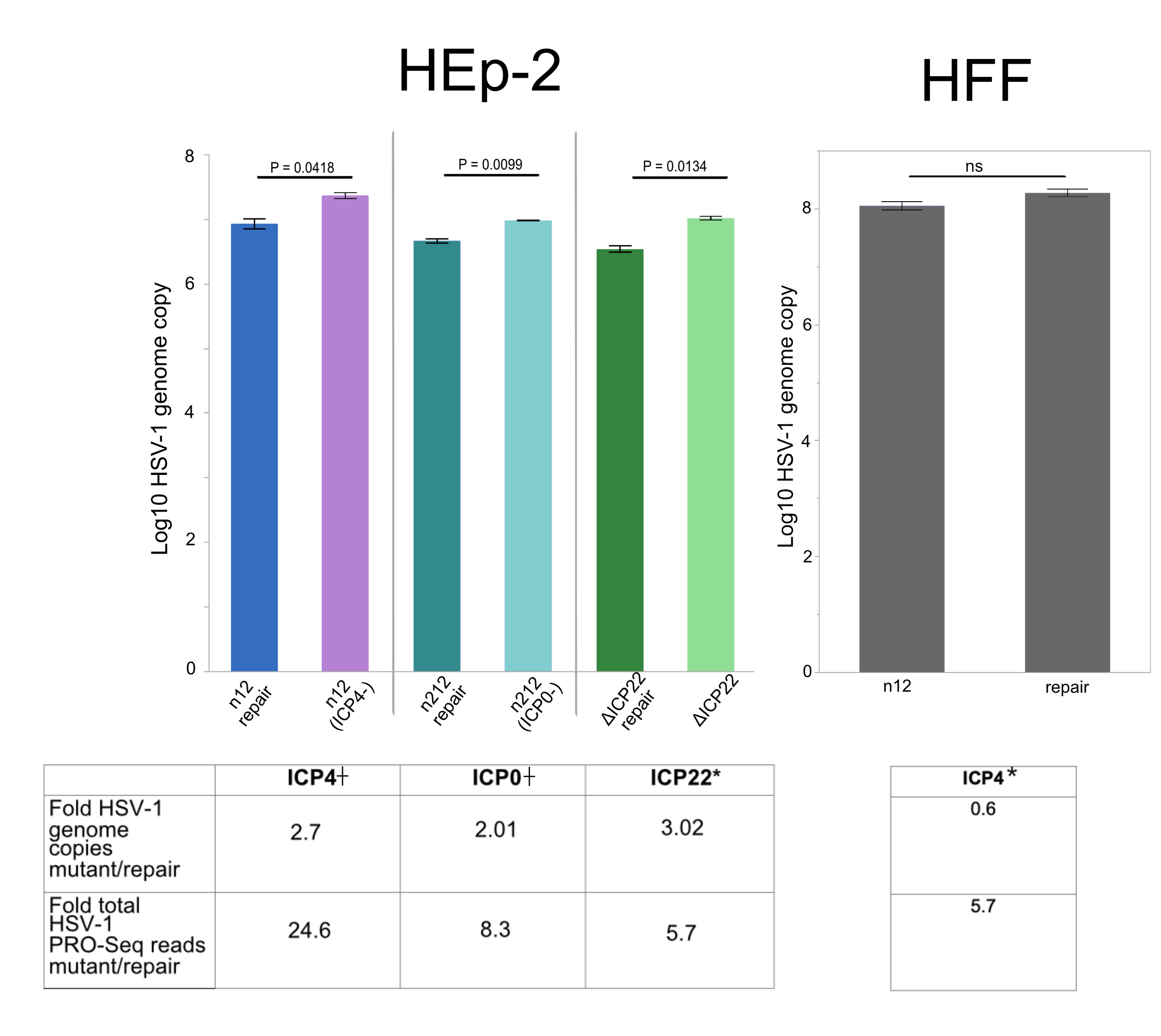
